# 2019-20 Wuhan coronavirus outbreak: Intense surveillance is vital for preventing sustained transmission in new locations

**DOI:** 10.1101/2020.01.24.919159

**Authors:** R.N. Thompson

## Abstract

The outbreak of pneumonia originating in Wuhan, China, has generated 830 confirmed cases, including 26 deaths, as of 24 January 2020. The virus (2019-nCoV) has spread elsewhere in China and to other countries, including South Korea, Thailand, Japan and USA. Fortunately, there has not yet been evidence of sustained human-to-human transmission outside of China. Here we assess the risk of sustained transmission whenever the coronavirus arrives in other countries. Data describing the times from symptom onset to hospitalisation for 47 patients infected in the current outbreak are used to generate an estimate for the probability that an imported case is followed by sustained human-to-human transmission. Under the assumptions that the imported case is representative of the patients in China, and that the 2019-nCoV is similarly transmissible to the SARS coronavirus, the probability that an imported case is followed by sustained human-to-human transmission is 0.37. However, if the mean time from symptom onset to hospitalisation can be halved by intense surveillance, then the probability that an imported case leads to sustained transmission is only 0.005. This emphasises the importance of current surveillance efforts in countries around the world, to ensure that the ongoing outbreak will not become a large global epidemic.

## 1. INTRODUCTION

The infectious agent driving the ongoing pneumonia outbreak (the 2019-nCoV) appears to have transitioned from animals into humans at Huanan seafood wholesale market in Wuhan, China [1–5]. Since then, cases have been recorded in other countries, and initial estimates suggest a case fatality rate of around 14% [6]. Even countries without confirmed cases are on high alert. For example, the United Kingdom has not yet seen a confirmed case, but officials are reported to be attempting to trace as many as 2,000 visitors that have travelled to that country from Wuhan.

The most devastating infectious disease outbreaks are those that have a wide geographical distribution, as opposed to being confined to a small region [7,8]. The previously known virus that is most similar to the 2019-nCoV is the SARS coronavirus [9], which generated cases in 37 countries in 2002-03 [9,10]. Since the 2019-nCoV is clearly capable of being transmitted by infected hosts to countries around the world, an important question for policy makers is whether or not these imported cases have the potential to generate sustained human-to-human transmission in new locations.

Here, we present data describing the times from symptom onset to hospitalisation for 47 patients from the current outbreak, obtained from publicly available line lists [11]. We fit an exponential distribution to these data, accounting for uncertainty due to the limited numbers of patients from whom data were available. Assuming that this distribution characterises the time spent by infected hosts generating new transmissions in the community, it is then possible to infer the probability that a case arriving in a new location is followed by an outbreak driven by sustained human-to-human transmission. We estimate this probability under the assumption that the transmissibility of the 2019-nCoV is similar to that of the SARS coronavirus, and then go on to consider the effect of shortening the mean time from symptom onset to hospitalisation. This provides an estimate of the risk that imported cases generate sustained outbreaks in new locations under different surveillance levels.

## 2. METHODS

### Time from symptom onset to hospitalisation

The distribution of times from symptom onset to hospitalisation was estimated using patient data from the ongoing outbreak [11] (data are shown in Fig 1A). Since the precise times of symptom onset and hospitalisation on the dates concerned were unknown, we converted the times from symptom onset to hospitalisation to intervals describing possible time periods. For example, for a case showing symptoms on 9 January 2020, and then being hospitalised on 14 January 2020, the time between symptom onset and hospitalisation lies between four and six days (see e.g. [12] for a similar calculation).

**Figure 1.**
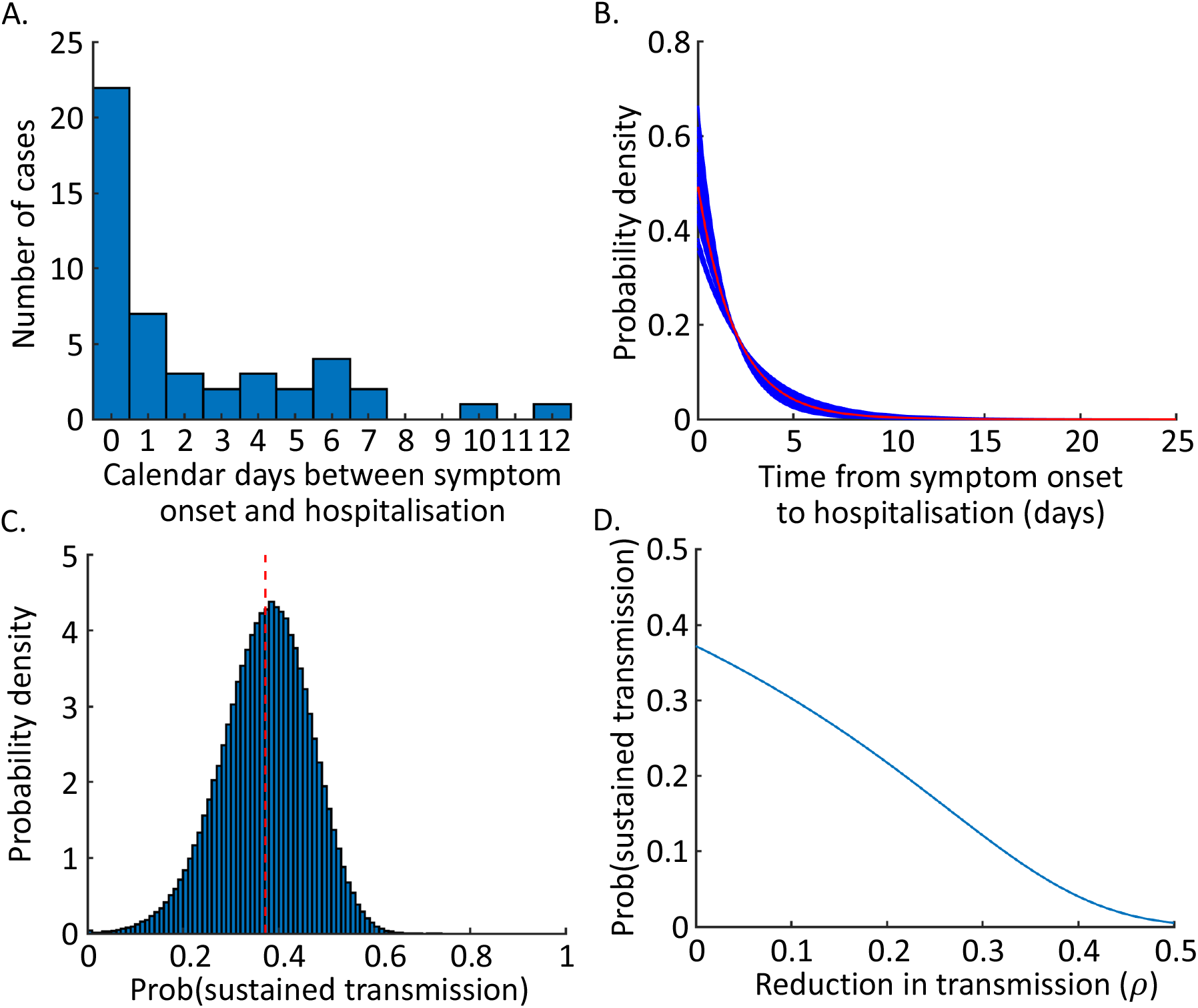
The probability of an outbreak driven by sustained human-to-human transmission arising following the importation one infected individual. A. Data describing the number of days between symptom onset and hospitalisation for 47 patients in the ongoing outbreak [11]. B. The estimated distribution of times between symptom onset and hospitalisation, estimated by fitting to the data shown in panel A. Blue lines show a range of equally possible distributions (see Methods; 50 distributions are shown here, selected at random from the *n* = 1,000,000 distributions considered), and the red line shows the average of the *n* = 1,000,000 distributions. C. The probability of sustained transmission for each possible distribution of times from symptom onset to hospitalisation (equation (1); blue histogram) and the probability of sustained transmission obtained by integrating over the possible distributions (equation (2); red line). D. The probability that a single imported case leads to sustained transmission in a new location, for different surveillance levels.

We then fitted the parameter (*γ*) of an exponential distribution to these interval-censored data using Markov chain Monte Carlo (MCMC). A chain of length 100,000,000 in addition to a burn-in of 100,000 was used. The chain was then sampled every 100 steps, giving rise to a range of *n* = 1,000,000 equally possible distributions for the times from symptom onset to hospitalisation, each characterised by a parameter estimate *γ_i_* (*i* = 1,2,..., *n*).

### Estimating the probability of sustained transmission

The distributions of times from symptom onset to hospitalisation were used to estimate the probability that an imported case will lead to sustained transmission, by assuming that infections occur according to a branching process (e.g. [13–15]). In this branching process, the effective reproduction number (accounting for control interventions, other than intensified surveillance which we model explicitly) of the 2019-nCoV when the virus arrives in a new location is denoted by *R* = *β/γ*, where the parameter *β* represents pathogen transmissibility [16]. We assumed that the transmissibility of the virus is similar to that of the SARS coronavirus, i.e. *β* = *R_SARS_ γ_SARS_*, where *R_SARS_* = 3 [17] and the mean infection duration for SARS is 1/*γ_SARS_* = 3.8 days [18].

The probability of a major outbreak [15,16] can be estimated for each of the equally possible distributions for the time from symptom onset to hospitalisation,

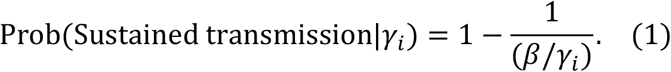

This can then be combined into a single estimate for the probability that an imported case leads to sustained transmission, *p*, given by

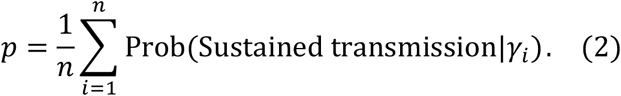

To include intensified surveillance in these estimates, we simply replaced the mean time from symptom onset to hospitalisation for each of the equally plausible distributions, 1/*γ_i_*, by the modified expression (1 - *ρ*)/*γ_i_*. In this approximation, the parameter *ρ* represents the reduction in the mean infectious period due to intensified surveillance.

### Multiple imported cases

The risk of sustained transmission given multiple imported cases was calculated by considering the possibility that none of those cases led to sustained transmission. Consequently,

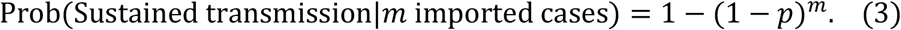

## 3. RESULTS

As described in Methods, the distribution of times between symptom onset and hospitalisation was estimated using Markov chain Monte Carlo (Fig 1B) from the patient data in Fig 1A. This gave rise to a range of equally plausible distributions describing these time periods (blue lines in Fig 1B). The average of these distributions is shown by the red line in Fig 1B, however we used the full range of distributions in our calculations of the probability of sustained transmission occurring from each imported case.

Each of the range of plausible distributions corresponded to an estimate for the probability of a major epidemic (equation (1) and histogram in Fig 1C). However, the probability of sustained transmission in fact takes a single value, which can be estimated by summing over the range of distributions using equation (2). The resulting probability of sustained transmission is 0.37 (red line in Fig 1C).

We then considered the reduction in the probability that an imported case leads to sustained transmission if surveillance is more intense. Specifically, we assumed that intensified surveillance led to a reduction in the mean period from symptom onset to hospitalisation, governed by the parameter *ρ* (where *ρ* = 0 corresponds to no intensification of surveillance, and *ρ* = 1 corresponds to an implausible scenario in which symptomatic cases are hospitalised immediately). We found that, if surveillance is intensified so that the mean time from symptom onset to hospitalisation is halved, the probability that each imported case leads to sustained transmission is reduced to only 0.005 (Fig 1D).

Finally, we considered the combined effect if multiple cases arrive in a new location. In that scenario, intense surveillance has the potential to significantly reduce the risk of sustained transmission compared to weak surveillance. For *ρ* = 0.5, the probability that any of 10 imported cases generate a substantial outbreak is only 0.049 (Fig 2C). This highlights the importance of rigorous surveillance, particularly in locations where infected hosts are most likely to travel.

**Figure 2.**
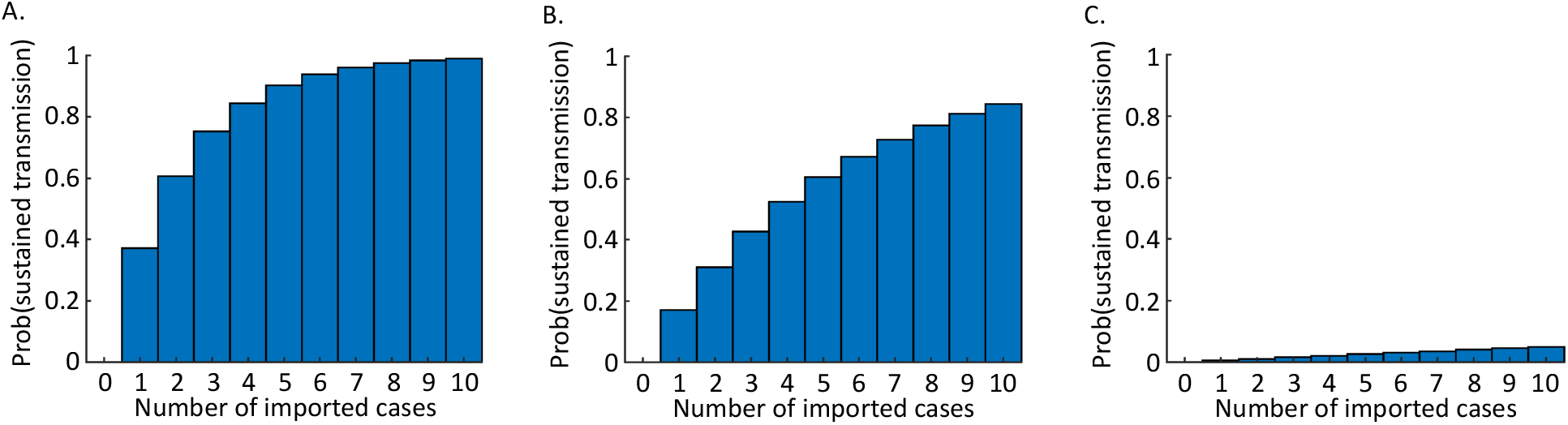
The probability of an outbreak driven by sustained human-to-human transmission arising from multiple imported cases, under different surveillance levels. A. No intensification of surveillance (*ρ* = 0). B. Medium level of surveillance intensification (*ρ* = 0.25). C. High level of surveillance intensification (*ρ* = 0.5). The results shown were calculated using equation (3).

## 4. DISCUSSION

There are concerns that the ongoing outbreak driven by 2019-nCoV could spread globally [3,5,19,20] with sustained transmission in countries around the world. In the near future, Chinese New Year presents a significant challenge, since this period often involves high travel rates, potentially leading to importations of the virus to many new locations [3,9].

Here, we have estimated the potential for cases arriving in new locations to lead to sustained transmission. According to the basic model that we have constructed, if surveillance levels are similar to those in China at the beginning of the current outbreak, and if this virus is similarly transmissible to the SARS coronavirus that spread globally in 2002-03, then the probability that each imported infected case generates an outbreak due to sustained transmission is approximately 0.37 (Fig 1C). However, under a higher level of surveillance, the risk of sustained outbreaks is substantially lower (Fig 1D). This result is particularly striking when multiple cases travel to a new location, either simultaneously or in sequence (Fig 2). In that scenario, intensified surveillance is particularly important.

Our study involves the simplest possible model that permits the risk of sustained transmission to be estimated from the very limited data that have been collected in this outbreak until now. As additional information becomes available, for example data describing virus transmissibility, then it will be possible to estimate the risk of outbreaks in new locations with more precision. We made the assumption that symptom appearance coincides with the onset of infectiousness. One of the features of the SARS outbreak in 2002-03 that allowed it to eventually be brought under control was the low proportion of onward transmissions occurring either prior to symptoms or from asymptomatic infectious hosts [21]. It might be hoped that infections due to 2019-nCoV share this characteristic.

Since our results were obtained using patient data from early in the ongoing outbreak, surveillance systems may not have been fully established when these data were collected, and patients may not have been primed to respond quickly to early symptoms. Our results might therefore be viewed as an upper bound on the risk posed by the 2019-nCoV. As the outbreak continues, it might be expected that the time from symptom onset to hospitalisation will decrease, leading to lower risks of sustained transmission, as has been observed for outbreaks of other diseases (e.g. the ongoing outbreak of Ebola virus disease in the Democratic Republic of the Congo).

Going forwards, it will be possible to include additional realism in the model. One example might be to consider spatial variation in host population densities and surveillance levels, leading to spatially inhomogeneous outbreak risks. It would also be possible to differentiate between mild and severe cases in the model. On the one hand, a mild case might be more likely to go unnoticed than a severe case, potentially leading to a higher outbreak risk. On the other hand, mild infections may generate fewer secondary cases than severe infections, thereby decreasing the outbreak risk. Complex interactions may therefore affect the risk of sustained transmission in an unpredictable fashion.

Despite the necessary simplifications made in this study, our analyses are sufficient to demonstrate the key principle that rigorous surveillance is important to minimise the risk of the 2019-nCoV generating large outbreaks in countries worldwide. We therefore support the ongoing work of the World Health Organization and policy makers from around the world, who are working with researchers and public health experts to manage this outbreak [2]. We also applaud efforts to make data publicly available [11]. Careful analysis of the outbreak, and minimisation of transmission risk as much as possible, are of clear public health importance.

## SUPPLEMENTARY MATERIAL

Data S1. The number of calendar days between symptom onset and hospitalisation for 47 patients from the ongoing pneumonia outbreak in Wuhan, China.

## COMPETING INTERESTS

There are no competing interests.

## FUNDING

This research was funded by Christ Church, Oxford, via a Junior Research Fellowship.

